# Microfluidic platform enables shear-less aerosolization of lipid nanoparticles for messenger RNA inhalation

**DOI:** 10.1101/2024.01.17.576136

**Authors:** Jeonghwan Kim, Antony Jozic, Elissa Bloom, Brian Jones, Michael Marra, Namratha Turuvekere Vittala Murthy, Yulia Eygeris, Gaurav Sahay

## Abstract

Leveraging the extensive surface area of the lungs for gene therapy, inhalation route offers distinct advantages for delivery. Clinical nebulizers that employ vibrating mesh technology are the standard choice for converting liquid medicines into aerosols. However, they have limitations when it comes to delivering mRNA through inhalation, including severe damage to nanoparticles due to shearing forces. Here, we introduce a novel <u>microfluidic aerosolization platform</u> (MAP) that preserves the structural and physicochemical integrity of lipid nanoparticles, enabling safe and efficient mRNA delivery to the respiratory system. Our results demonstrated the superiority of the novel MAP over the conventional vibrating mesh nebulizer, as it avoided problems such as particle aggregation, loss of mRNA encapsulation, and deformation of nanoparticle morphology. Notably, aerosolized nanoparticles generated by the microfluidic device led to enhanced transfection efficiency across various cell lines. *In vivo* experiments with mice that inhaled these aerosolized nanoparticles revealed successful, lung-specific mRNA transfection without observable signs of toxicity. This pioneering MAP represents a significant advancement for the pulmonary gene therapy, enabling precise and effective delivery of aerosolized nanoparticles.

## INTRODUCTION

The successful development of mRNA vaccines against SARS-CoV-2 has transformed how the researchers and clinicians perceive the use of mRNA as a therapeutic tool^1^. Previously considered a theoretical approach, it has now become a promising and feasible option for clinical applications^2, 3^. Particularly, mRNA has garnered significant attention in the field of pulmonology for its potential in treating inherited diseases^4, 5^, including cystic fibrosis (CF) and alpha-1 antitrypsin deficiency as well as in the field of vaccinology for intranasal vaccination approaches^6^. Accordingly, research exploring mRNA therapy for the pulmonary system has experienced rapid growth^7, 8^. Nevertheless, the most significant challenge in its clinical translation lies in identifying a safe and effective delivery strategy^8^. The recent progress in the development of novel nanoparticles capable of targeting the lungs following systemic administration shows great potential for lung-specific mRNA delivery^9,10^. However, concerns regarding the potential risk of off-target delivery persist with these RNA vectors. Given that many pulmonary diseases are closely linked to conditions in the pulmonary epithelial cells, the most direct, safe, and efficient method for administering mRNA therapies might be via inhalation^8^. Inhalation allows for the therapy to reach high concentrations in the pulmonary epithelial tissues, enabling targeted and localized treatment for respiratory conditions while minimizing systemic exposure. This local administration holds significant promise for the treatment of the pulmonary system using mRNA therapies. Clinical trials based on inhalation-mediated mRNA delivery (NCT05712538 and NCT05737485) indicate the growing interest in this approach. However, inhaled mRNA therapy continues to encounter challenges related to the shearing damage caused by the inhalers, particularly nebulizers, which are commonly used to generate aerosols from aqueous medications^4, 11^. Both jet nebulizers and vibrating mesh nebulizers involve an atomization process that exerts strong shearing forces on the nanoparticles, leading to a significant loss in their mRNA delivery efficiency. Thus, addressing and mitigating these challenges are critical to maximize the efficacy and safety of inhaled mRNA therapies for pulmonary diseases^8^. Studies focused on optimizing nanoparticles have shown promising results in improving their ability to endure the shearing forces exerted by nebulizers^12–14^. Research conducted by our group and others previously demonstrated that increasing the molarity of polyethylene glycol (PEG) in lipid nanoparticle (LNP) formulations enhanced mRNA transfection after nebulization^4, 15^. High PEG contents in LNPs could improve their resilience to damage during the aerosolization process, attenuating the negative impact of shear forces on mRNA delivery efficiency.

While fine-tuning nanoparticles may rescue their delivery competence during nebulization, an alternative solution to this challenge lies in the development of novel aerosolization platform^16^. Nebulization through a vibrating mesh essentially involves directing fluid flow through a fine screen to atomize aerosols on a microscale. In this context, exploring other possible fluid mechanics for plume generation presents a promising chance to develop innovative aerosolization devices. Microfluidic devices have been widely employed in the production of various nanoparticles, providing continuous, controllable, and reproducible manufacturing of small-sized nanoparticles with a narrow size distribution in a simple process^17–19^. We hypothesized that utilizing microfluidics to flow LNPs through micro-channels offers the potential for aerosolizing mRNA without subjecting the nanoparticles to damaging shearing forces.

In this study, we have developed a novel, translatable, microfluidic-based aerosolization platform (MAP) that generates uniform aerosols containing mRNA encapsulated within LNPs (LNP/mRNA), effectively circumventing damaging shear forces. A microfluidic chip is connected to a cartridge that contains LNP/mRNA solution, providing on-demand droplet generation with precise dose control capabilities potentially for either individual chronic treatment or mass vaccination purposes. Unlike the ultrasonication method in vibrating mesh nebulizers or the jet explosion method in jet nebulizers, droplet generation process in this MAP expels droplets with minimal shear-affected volume, making it highly suitable for delivering macromolecule-based therapies, such as nucleic acids, proteins, and nanoparticles. We compared our MAP to the clinical-grade vibrating mesh nebulizer based on ultrasonication method for droplet generation. For the studies, we used a gold standard ionizable lipid with formulations previously optimized for siRNA and mRNA delivery to highlight clinical translation of our MAP. This innovative platform holds promise to address the clinical need of inhaled nanoparticles for a broad range of RNA therapeutics and vaccines by offering a controlled and efficient aerosolization without compromising the integrity of the nanoparticles.

## RESULTS

To achieve effective delivery of LNP-assisted gene therapies in the form of micron-sized droplets, we utilized a microfluidic device consisting of a complementary metal oxide semiconductor (CMOS) based chip with integrated microfluidic structures. This device comprises an array of 960 droplet ejectors, each individually addressable, enabling the generation of droplet plumes containing LNP/mRNA (Fig. 1A). Briefly, the microfluidic chip is monolithically fabricated on a silicon base layer through standard photolithographic techniques, incorporating a polymeric manifold and nozzle configuration. It is a specialized device that utilizes microfluidic and thermodynamic principles to flow the liquid through the fluid manifolds and to generate and eject small droplets of liquid. Each of the 960 droplet ejectors consists of a microfluidic chamber that holds the liquid, and a nozzle or orifice (approximately 10 µm in diameter) through which the liquid droplets are expelled (Fig. 1B, C). Underneath the chamber, multiple thin film layers of heater elements and electric materials are integrated. The device is actuated digitally by applying short electrical pulses to individual heater elements, generating microbubbles that propel the droplets out of the nozzle. In detail, when an electric current is passed through the resistive heating element, the temperature of the surface of the heater increases rapidly, causing the liquid in substantial contact with the heater surface to vaporize. As the liquid vaporizes, it forms a bubble within the microfluidic channel, and the sudden expansion of this bubble propels a droplet out through the nozzle (Fig. 1D). Proper control of the electrical pulse duration and magnitude consistently generates droplets of a specified size and speed, while actuation of individual ejectors allows for precise and controlled dispensing. For example, if each of the 960 ejectors are actuated individually and repeatedly at a frequency of 10 kHz per nozzle, droplets are produced at a rate of 9.6 million droplets per second, resulting in the production of a plume (Supplementary Fig. 1 and Supplementary Video 1). Total dispensed volume is adjustable and easily controlled by specifying the exact number of droplets to be ejected per nozzle. During the entire procedure, the liquid is precisely controlled and conveyed to the nozzle through microfluidic techniques, ensuring a consistent and uniform droplet size and shape. Furthermore, the microfluidic techniques enable accurate control over the plume dimensions, droplet velocity and size, as well as the administered dose.

**Fig. 1.**
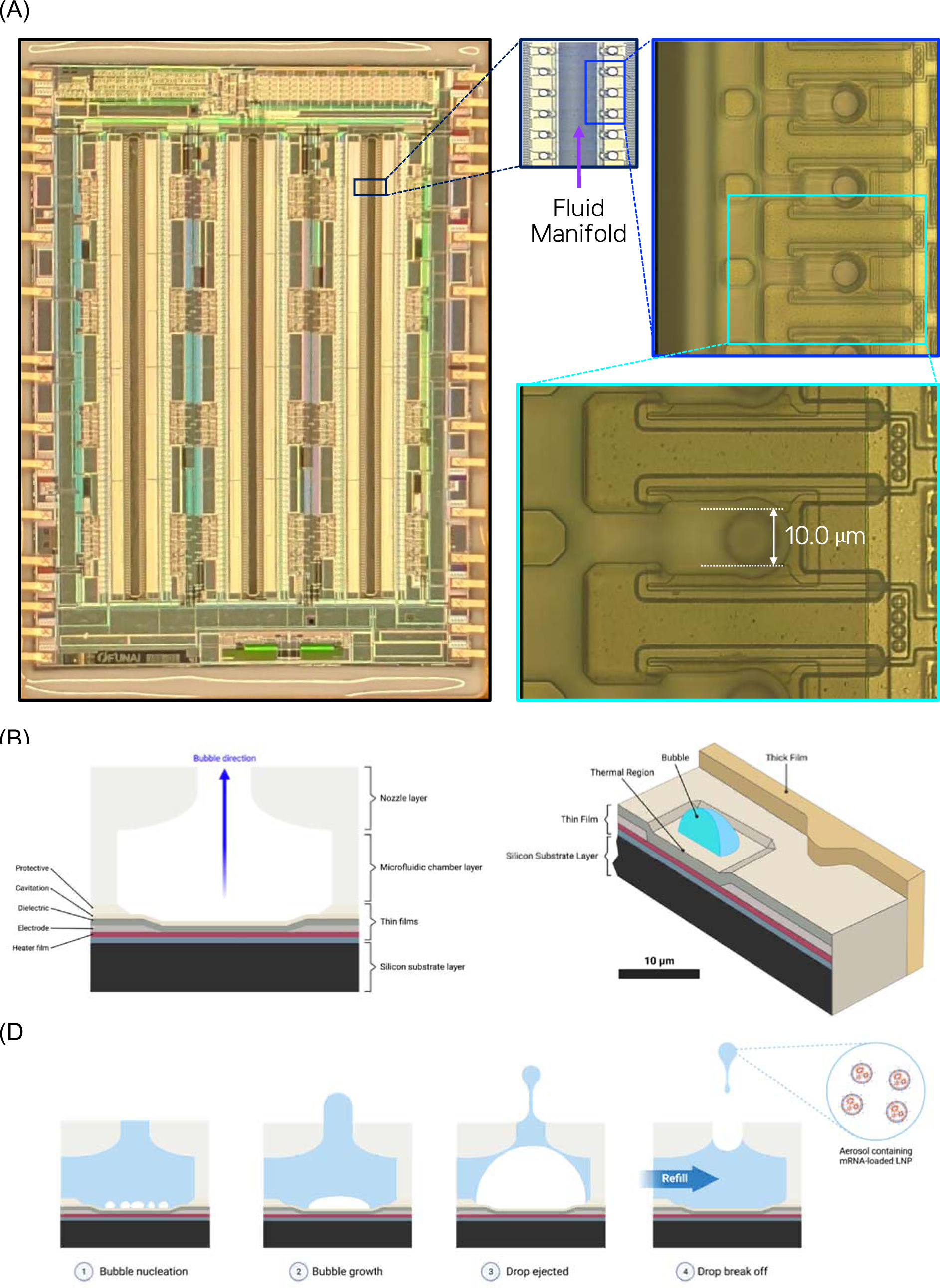
Microfluidic system to aerosolize LNP/mRNA. (A) Microscopic images of a microfluidic chip cartridge containing 960 nozzles. The nozzles are arrayed into three fluid channels, each channel consisting of 2 columns of nozzles and each channel linked to separate fluid chambers. The diameter of each nozzle is approximately 10 µm. (B) Illustration of the cross-sectional view of an individual nozzle made using a conventional semiconductor CMOS process, proprietary FMS thin film processes, and proprietary FMS MEMS photolithography processes. The thin film consists of various layers, including a protective layer, cavitation layer, dielectric film, electrode, and heater film. Blue arrow indicates primary direction of bubble growth when the heater is actuated. (C) Illustration of the layer structure of the microfluidic cartridge head with a bubble. Scale bar indicates 10 µm. (D) Schematic illustration explaining the sequential mechanism of ejection (aerosolization) of LNP/mRNA droplets at the cross-section of a nozzle: (1) bubble nucleation, (2) bubble growth, (3) drop ejected, and (4) drop break off and refill.

In inhalation-mediated drug delivery, the droplet size plays a pivotal role in determining its site of accumulation within the pulmonary tissues. Larger droplets tend to deposit in the upper airways and are susceptible to removal through mucociliary clearance before reaching the deeper lung tissues. Conversely, smaller droplets have a higher chance of penetrating deeper into the lungs and reaching the alveolar region. Therefore, the ideal droplet size for inhalation-mediated drug delivery depends on the specific target site of action. For conditions like asthma, chronic obstructive pulmonary disease (COPD), or CF, smaller droplets capable of reaching the lower airways and alveoli are preferred. On the other hand, larger droplets may be more suitable for topical treatment of upper respiratory infections or inhaled vaccinations. Hence, precise control over droplet size in aerosols is vital for the successful development of effective delivery platforms for inhalation-mediated medication. To assess the droplet size generated by our MAP, we conducted analyses of their size distribution using two different techniques: a JetXpert drop watcher and a Spraytec^®^ droplet size measurement system. For the droplet size characterization, we used liposomes (DSPC:cholesterol:DSPE-PEG2K = 52:45:3) to mimic physical properties of bulk LNP suspensions. The mean size of the liposomes was approximately 80 nm with an approximate polydispersity index (PDI) 0.1 (Supplementary Fig. 2). The drop watcher system allows us to capture precise images of individual droplets in flight, measure their shape, and estimate their volume, even at the pico-liter scale. We compared drop formation of droplets ejected from the nozzles of the microfluidic cartridge at two different operating frequencies, resulting in each nozzle being actuated at 1 kHz and 15 kHz, respectively, for visualization of a single drop ejection event (Fig. 2A-B). Drops generated at 1 kHz were observed to break apart into 4 droplets in flight (Fig. 2A), while those at 15 kHz broke up into 3 droplets (Fig. 2B). By analyzing the captured images of the ejected fluid upon exiting the nozzle opening, it was revealed that 4 droplets formed at 1 kHz ejection frequency contained 59%, 30%, 8%, and 3% of the total ejected volume, respectively (Fig. 2C). Similarly, when operating at 15 kHz frequency, the three droplets formed per ejection were 72%, 17%, and 11% of the total ejected volume (Fig. 2D). It is noteworthy that, because the droplets have different velocities, they collide with each other, resulting in the breakup or merge in the air. As a result, the drop watcher system is limited to the analysis of the breakup of droplets upon ejection from the nozzle opening^20^. To accurately determine the droplet size of the aerosols, we employed Spraytec®, a size determination method based on laser diffraction of aerosols^21^. Droplet sizes were measured at two ejection frequencies – at 7.5 kHz and 15 kHz – using all of the nozzles (Fig. 2E, F). At 7.5 kHz, 50% of the total number of drops were less than 13.0 µm (Dn(50)) and 90% of the drops were less than 20.9 µm (Dn(90)) (Fig. 2E). Similarly, at 15 kHz, 50% of the total number of drops were less than 14.6 µm ((Dn(50)) and 90% of the drops were less than 24.4 µm (Dn(90)) (Fig. 2F). We also characterized the droplet size distribution as measurement parameter varied, including measuring distances, the number of nozzles, and the elapsed time of aerosolization. When the measurement was taken near the nozzle plate, the distribution became slightly narrower (Supplementary Fig. 3). It was observed that lowering the frequency of aerosolization or the number of nozzles tended to slightly decrease the droplet size (Supplementary Fig. 4). Additionally, we measured the size over the course of elapsed time during aerosolization. The size distribution was smaller at the beginning of aerosolization compared to the middle and end, suggesting that the aerosolization process becomes more stable as dispensing continues (Supplementary Fig. 5). It is noteworthy that the impact of the studied variables on droplet size was consistently less than 2 µm in all cases, indicating the robust aerosolization performance of the MAP (Supplementary Fig. 3-5).

**Fig. 2.**
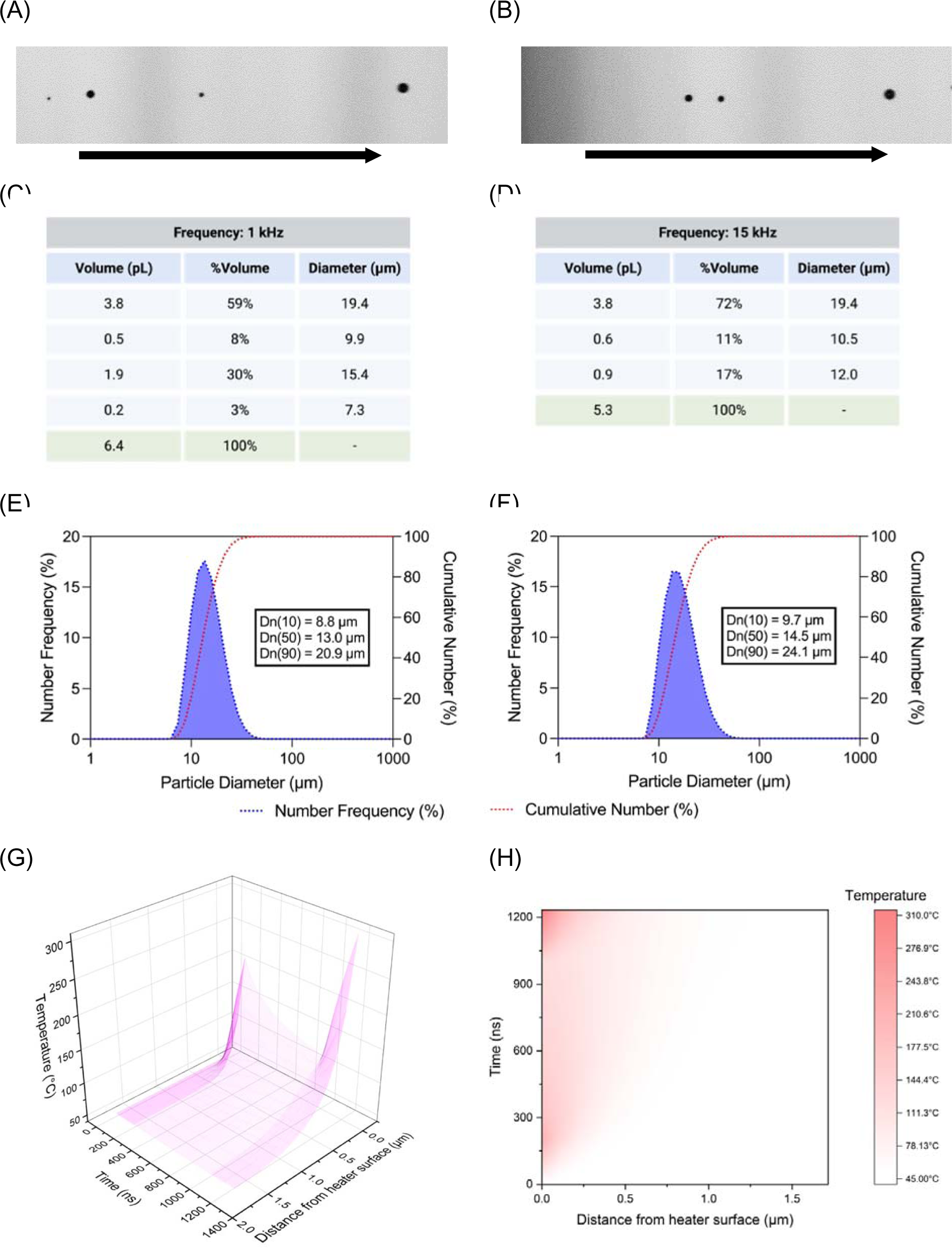
Characterization of droplets produced from the MAP. (A-F) Representative images of droplets from a single ejection at (A) 1 kHz and (B) 15 kHz frequencies. Black arrow indicates the direction of plumes. The volumetric percentage of each droplet was estimated from the captured images and recorded for each ejection at (C) 1 kHz and (D) 15 kHz. (E, F) The laser diffraction-based determination of the size distribution of droplets ejected at frequencies of (E) 7.5 kHz and (F) 15 kHz. (G,H) Electric-thermal simulation of variation of fluid temperature in thermal boundary layer during pulse according to the geometric dimension and time. (G) Response surface plot and (H) contour plot of fluid temperature during pulse. Time is presented as the duration from the start of pulse in nanoseconds (ns).

Analysis was also performed, via simulation to estimate the temperature change of the fluid in the thermal boundary layer during the electric pulse, considering the effects of time and geometric dimensions. The temperature distribution of the fluid in the ejector chamber was non-uniform, with higher temperatures closer to the heater surface (Fig. 2G, H). As the distance from the heater surface exceeded 1 µm, the temperature increase was minimal. Additionally, the temperature rise was highly correlated in time with the actuation of the drop ejection with the electrical signals to the heater, with the peak temperature near the end of the ejection sequence (Fig. 2G, H). Although the temperature experienced a localized increase to 300°C for only a few microseconds, this rise was limited to the regions in proximity to the heater, which is unlikely to compromise the overall stability of the LNP/mRNA to be aerosolized.

To examine any potential impact of thermal changes and shearing forces generated by the MAP, we attempted to aerosolize LNPs and deliver encapsulated mRNA to the cells. The LNP formulation consisted of DLin-MC3-DMA, cholesterol, DSPC, and DMG-PEG_2000_ at a molar ratio of 50:38.5:10:1.5 (i.e., the lipid composition of patisiran), with the mRNA present at an N/P ratio of 5.30 (Fig. 3A). Following the aerosolization of LNP/mRNA using either a vibrating mesh nebulizer or our MAP, we compared the hydrodynamic size distributions, zeta-potentials, and RNA encapsulation of the nanoparticles with those of the untreated nanoparticles (Fig. 3B-G). Untreated LNP/mRNA were < 100 nm in hydrodynamic diameter with narrow distributions (PDI < 0.2) (Fig. 3B-D). When aerosolized by a vibrating mesh nebulizer, the size distribution of the nanoparticles was changed dramatically. As shown previously^4, 11^, the ultrasonication of a mesh nebulizer resulted in large, unstable, and aggregated LNPs. The nebulized LNP/mRNA displayed approximately 700 nm hydrodynamic diameter with wide distributions (PDI > 0.7) (Fig. 2B-D). In contrast, the size distributions of the nanoparticles were scarcely affected by the MAP (Fig. 3B-D). The hydrodynamic sizes of the nanoparticles that went through microfluidic aerosolization were < 100 nm with narrow distribution (PDI < 0.2) (Fig. 3C, D). Despite slight decreases after aerosolization, zeta-potentials of all LNP samples were in the neutral range (i.e., between +10 and −10 mV) (Fig. 3E). These conspicuous differences in the dynamic light scattering (DLS) analysis indicate that the MAP provides clear benefits in retaining the physicochemical properties of LNP/mRNA during aerosolization than traditional mesh nebulizers. We further characterized the encapsulation state of mRNA after LNP aerosolization. While untreated LNP/mRNA had high encapsulation (> 95%), the ultrasonication of the mesh nebulizers led to significant loss of mRNA encapsulation (ca. 43%) (Fig. 3F). Again, the microfluidic aerosolization rescued the encapsulated mRNA from the leakage, indicating its protective effects of mRNA cargos during aerosolization. To confirm the retainment of cargo encapsulation, the LNP/mRNA samples were subject to the agarose gel electrophoresis analysis (Fig. 3G, Supplementary Fig. 6). mRNA solution (‘mRNA only’ group) and the LNP/mRNA treated with Triton X-100 detergent showed mRNA migration in the gel, generating two bands: one for its linear form and the other for its secondary structure. Untreated LNP/mRNA did not migrate but remain in the wells, indicating the stable encapsulation of mRNA within the nanoparticles. When tested the LNP/mRNA exposed to the vibrating mesh nebulization, the mRNA bands appeared, indicating that the nanoparticles leaked the mRNA during nebulization (Fig. 3G). By contrast, the ones aerosolized by the MAP displayed very faint bands in the gel, supporting the negligible amount of mRNA loss during aerosolization (Fig. 3G). These results showed that the MAP can aerosolize LNPs without the risk of mRNA loss (Fig. 3G), in conjunction with the results of mRNA encapsulation assay (Fig. 3F). We reasoned that the loss of mRNA during nebulization may arise from aggregation and rearrangement of lipids incorporated in the LNPs. To examine this hypothesis, we tested if the LNP/mRNA go through the membrane fusion in the nebulization processes through fluorescence resonance energy transfer (FRET)^22, 23^. Liposomes containing two DOPE-conjugated FRET probes, 7-nitrobenzo-2-oxa-1,3-diazole (NBD-PE) and lissamine rhodamine B (Rho-PE), were prepared, resulting in what we refer to as FRET liposomes (Fig. 3H). The proximate presence of both probes in the liposomes caused a decrease in NBD fluorescence due to FRET occurring between NBD and rhodamine B. Following the process of membrane fusion, the distance between the two probes increased, leading to a subsequent rise in NBD fluorescent signals. To assess membrane fusion events, LNP/mRNA were mixed with FRET liposomes and subjected to various conditions: Triton X-100, bath sonication, vibrating mesh nebulization, and MAP (Fig. 3I). The signal obtained from the untreated sample represented 0% fusion, while Triton X-100 treatment represented 100% fusion. Among the tested methods, vibrating mesh nebulization induced the highest degree of membrane fusion (Fig. 3I). Notably, it significantly surpassed bath sonication, likely due to the additional shear stresses generated by the pores of the mesh. On the other hand, microfluidic aerosolization resulted in significantly lower levels of fusion (approximately 16% lower), indicating its mild effects on the lipid membrane structure during LNP aerosolization. In summary, our MAP offers distinct advantages in the aerosolization of LNP/mRNA by preserving the physicochemical properties and structural integrity of the nanoparticles and providing better protection for the encapsulated mRNA cargos, surpassing the conventional vibrating mesh nebulizer in these aspects.

**Fig. 3.**
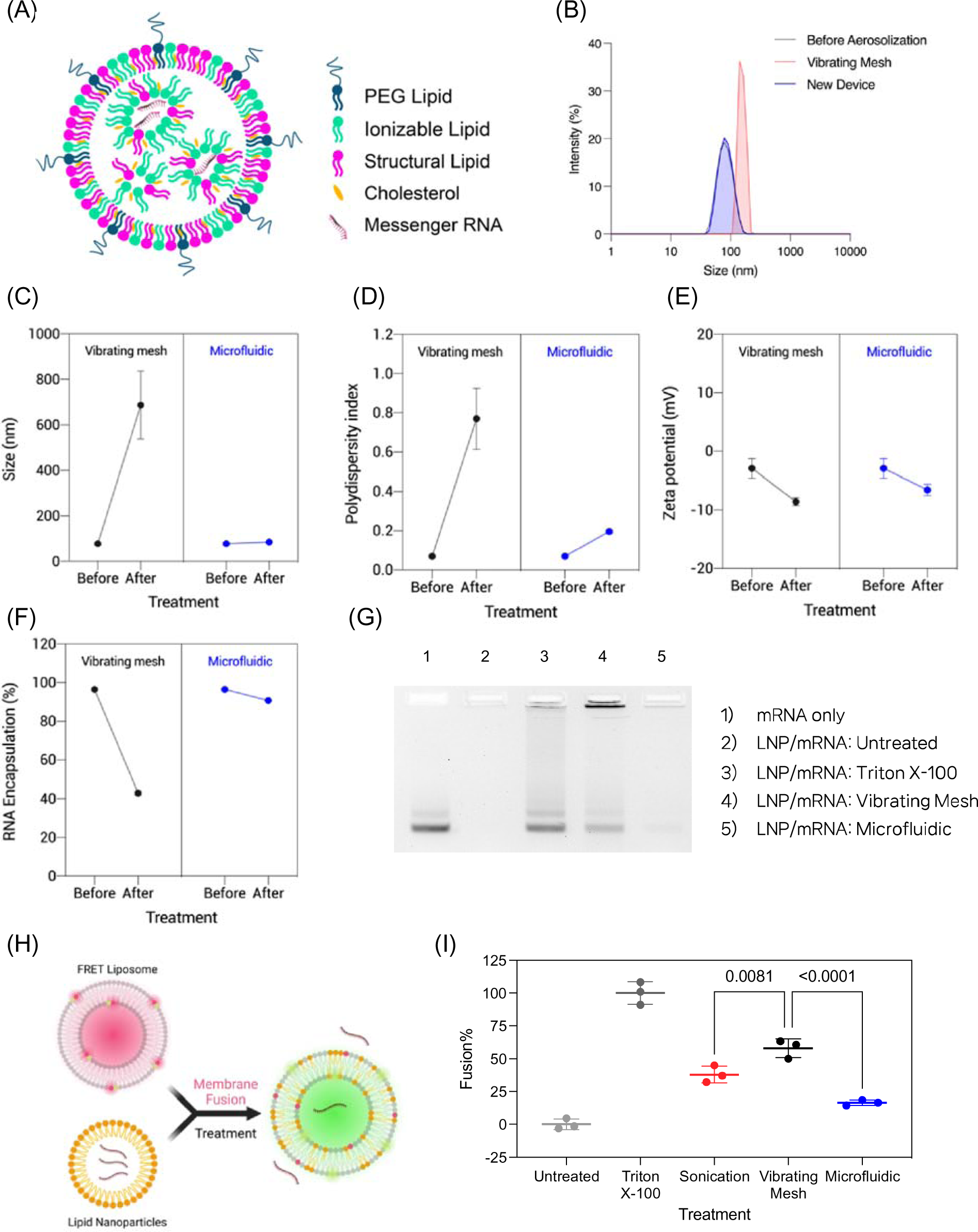
Physicochemical characterization of LNP/mRNA aerosols produced by a conventional nebulizer and the MAP. (A) A schematic representation of a single LNP/mRNA. PEG lipid (blue); ionizable lipid (green); structural lipid (pink); cholesterol (yellow). (B) Representative size distributions of LNP/mRNA for different treatments: no treatment (gray), vibrating mesh (red), and MAP (blue). (C-F) Changes in LNP/mRNA following aerosolization via a vibrating mesh (black) or MAP (blue): (C) size, (D) polydispersity index, (E) zeta potential (mV), and (F) mRNA encapsulation. (G) A representative image showing agarose gel electrophoresis analysis from various treatment conditions: 1) mRNA only, 2) untreated LNP/mRNA, 3) LNP/mRNA + Triton X-100, 4) LNP/mRNA + vibrating mesh, and 5) LNP/mRNA + MAP. (H) A schematic diagram of FRET-based lipid membrane fusion assay. (I) FRET-based assay results for lipid membrane fusion of LNP/mRNA exposed to different conditions.

Next, we examined the morphology of LNP/mRNA after undergoing aerosolization using cryogenic transmission electron microscopy (CryoTEM). The untreated LNP/mRNA sample displayed a spherical shape with a single bilayer (Fig. 4A), and the particle diameter was found to be less than 100 nm, consistent with the DLS analysis results (Fig. 2C). However, ultrasonication with mesh nebulizers caused the dissociation and aggregation of the nanoparticles (Fig. 4B). The density of nanoparticle per field of view was also notably reduced (Data not shown). Some particles captured exhibited diameters in the range of several hundred nanometers, and the images had a generally low signal-to-noise ratio, indicating poor electron density of the sample on the grid. In contrast, the morphology of LNP/mRNA following microfluidic aerosolization closely resembled that of the untreated sample, despite the presence of a few large nanoparticles (Fig. 4C). The macroscopic observation of the LNP/mRNA samples further supported that the ultrasonication processes led to the dissociation and aggregation of LNPs. Comparatively, the untreated sample and the one subject to microfluidic aerosolization appeared optically translucent. However, the sample aerosolized by the mesh nebulizer became opaque (Supplementary Fig. 7), suggesting the presence of large particles capable of scattering light.

**Fig. 4.**
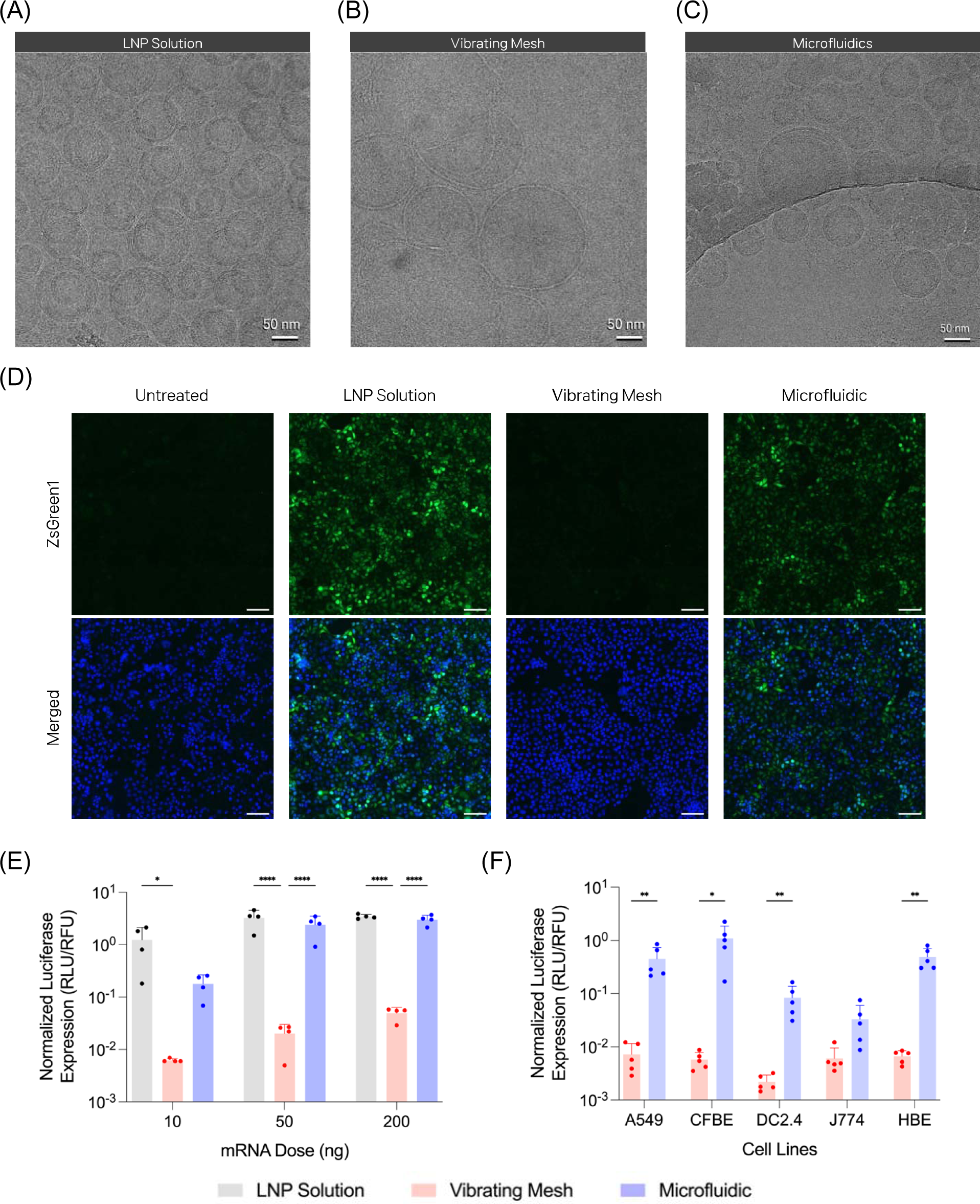
Performances of LNP/mRNA aerosols produced by the MAP surpasses that of a vibrating mesh nebulizer. (A-C) Cryogenic transmission electron microscopy (cryoTEM) images of LNP/mRNA (A) in solution, (B) after aerosolization by a vibrating mesh nebulizer, and (C) after aerosolization by the MAP. Scale bars indicate 50 nm. (D) Fluorescence microscopy images of HeLa cells subjected to various conditions: untreated, treated with LNP/ZsGreen1 in solution, aerosolized by a vibrating mesh, and aerosolized by the MAP (from left to right) at a dose of 1 µg mRNA per chamber. Green represents ZsGreen1 proteins and blue represents nuclei. Scale bars indicate 200 µm. (E) Normalized luciferase expression of 293T/17 cells treated with LNP/Fluc solution (light blue), LNP/Fluc aerosolized by a vibrating mesh nebulizer (red) or the MAP (blue) at various mRNA doses. (n=4) (F) Normalized luciferase expression of various cell lines by LNP/Fluc aerosolized by a vibrating mesh nebulizer (red) or the MAP (blue) at a dose of 50 ng mRNA per well. (n=5) *p<0.05; **p<0.01; ****p<0.0001.

We hypothesized that the intact nanostructure provided from MAP would produce greater mRNA transfection than the damaged nanostructure by mesh nebulizers. To validate this assumption, we treated 293T/17 cells with LNPs containing ZsGreen1 mRNA (LNP/ZsGreen1) with or without aerosolization. After 24 h incubation post-treatment, ZsGreen1 mRNA transfection was assessed using fluorescence microscopy. The negative control group showed no green fluorescence, while the cells treated with the LNP/ZsGreen1 solution exhibited bright green fluorescence across the field of view (Fig. 4D). When comparing the vibrating mesh nebulizer and the MAP, it became evident that the latter resulted in a greater expression of ZsGreen1 protein in the treated cells (Fig. 4D). Moreover, the level of ZsGreen1 expression observed in cells treated with aerosolized LNP/ZsGreen1 using the MAP looked comparable to that of cells treated with the LNP/ZsGreen1 solution. It suggests that the aerosolization of LNP/mRNA by the MAP has little impact on the efficiency of nanoparticle-mediated mRNA delivery to cells. On the other hand, vibrating mesh nebulizers noticeably hindered *in vitro* mRNA transfection by nanoparticles during aerosolization. We conducted additional measurements to examine the advantages of the MAP compared to vibrating mesh nebulizers in mRNA delivery. When delivering firefly luciferase (Fluc) mRNA instead, we found consistent patterns again. In 293T/17 cells, the group treated using the vibrating mesh nebulizer showed a nearly 100-fold reduction in mRNA transfection compared to the LNP solution treatment group (Fig. 4E). In contrast, the group treated with the MAP displayed mRNA transfection levels similar to the LNP solution treatment group, albeit an approximately 7-fold decrease observed only in the lowest mRNA dose (Fig. 4E). All treatments had little effect on the cell viability of 293T/17 cells (Supplementary Fig. 8). To corroborate the benefit of the MAP in aerosolizing LNP/mRNA, we iterated the experiments in more biologically relevant cells. In human bronchial epithelial cells (16HBE14o-), the MAP showed comparable efficiency in delivering mRNA to that of the LNP solution treatment (Supplementary Fig. 9). By contrast, the vibrating mesh nebulizer significantly impaired the nanoparticles’ ability to deliver mRNA across all tested mRNA doses. The subsequent screening across various cell lines exhibited similar results, demonstrating that the MAP achieved mRNA delivery efficiencies approximately 5 to 187 times higher than the vibrating mesh nebulizer (Fig. 4F, Supplementary Fig. 10). In sum, our MAP clearly demonstrated significant advantages over the vibrating mesh nebulizer, as it displayed superior efficiency in delivering mRNA to cells.

Having established the overall relative advantages of our MAP in generating nanoparticle-containing aerosols, we proceeded to deliver mRNA to the mouse lungs. For this purpose, we employed a whole-body rodent inhalation system, which consisted of a 3 L container connected with the MAP (Fig. 5A). This system facilitated the controlled administration of aerosolized LNP/mRNA to the mice, enabling the evaluation of its potential applicability for inhalable mRNA therapy. The MAP was programmed through the controller to generate a plume, and we loaded the cartridge with LNPs containing Nluc mRNA (LNP/Nluc). The mice were placed inside the chamber and exposed to the aerosol containing LNP/Nluc, after which they spontaneously inhaled the nanoparticles. To confirm the successful pulmonary delivery of mRNA to the lungs using the rodent inhalation system facilitated by our MAP, we conducted RNAScope *in situ* hybridization (ISH) analysis.

**Fig. 5.**
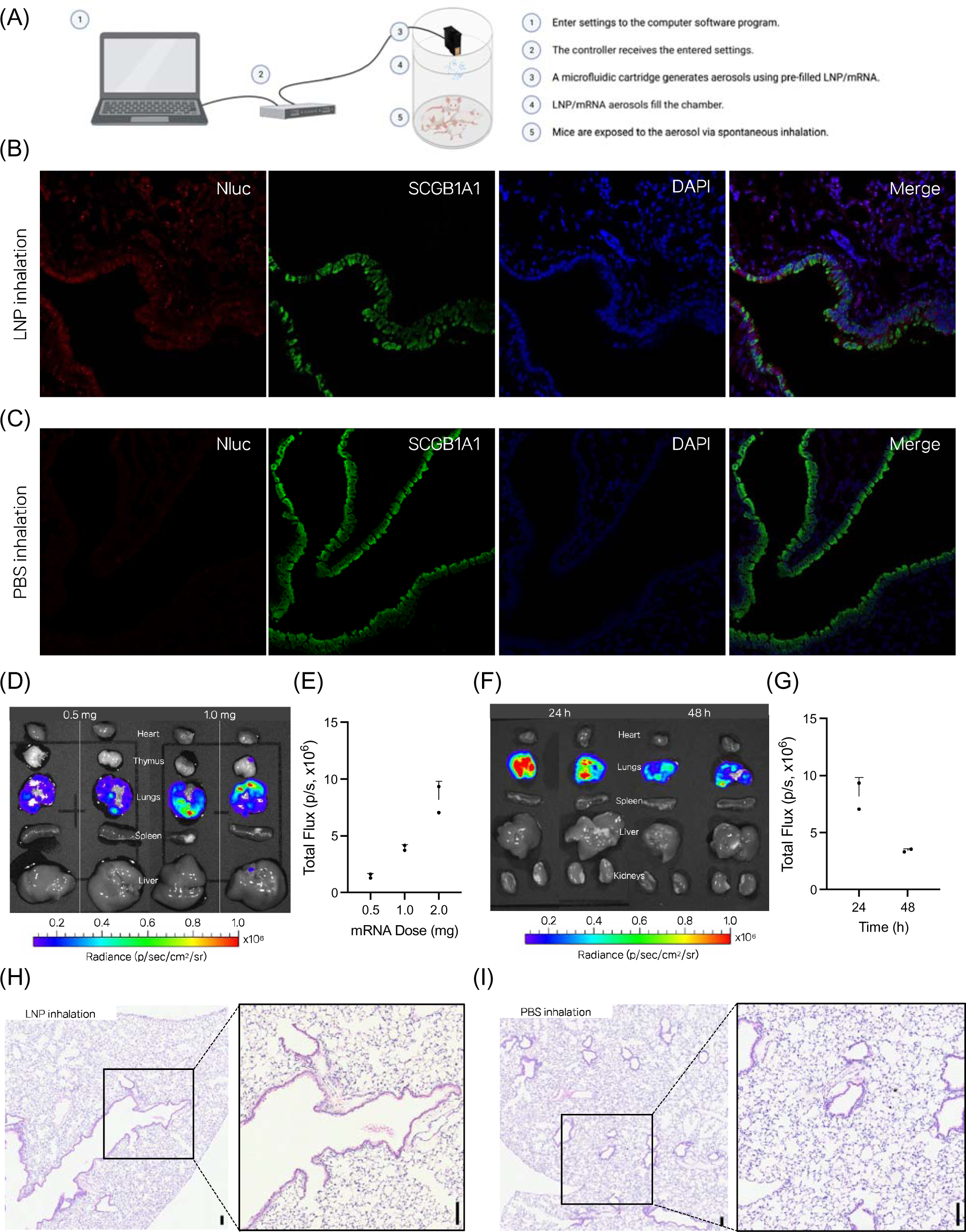
Effective mRNA delivery to the lungs using microfluidics-assisted aerosolization. (A) A schematic diagram of a whole-body rodent inhalation system, which consisted of a 3 L container connected with the MAP. (B,C) Representative images of RNAscope *in situ* hybridization analysis of BALB/c mouse lung sections after inhalation of (B) LNP/Nluc inhalation or (C) sterile PBS using the MAP. Nluc mRNA transcript (red), SCGB1A1 (green), and nuclei (blue). 20× magnification. (D-G) Ex vivo luminescence (D,F) images and quantification (E,G) in lungs after LNP/Nluc were aerosolized and administered to mice. (D,E) The variations included changing the amounts of mRNA to be aerosolized when the images were taken at 24 h post-inhalation (D,E), or conducting luminescence imaging at different time points when 2.0 mg of mRNA was aerosolized (F,G). (n=2) (H,I) Histopathological analysis of mouse lungs collected 24 h after (H) inhalation of LNP/Nluc when 1.0 mg of mRNA was aerosolized, or (I) inhalation of an equal volume of sterile PBS using the MAP. Scale bars indicate 100 µm.

We aerosolized LNP/Nluc containing 1 mg of mRNA into the chamber and allowed the mice to inhale the aerosols. After 24 hours, we collected the mouse lungs, fixed them, and prepared sections for staining. Using confocal imaging, we visualized the presence of Nluc mRNA transcripts in the lungs. Additionally, we labeled club cells, a specific type of bronchial epithelial cells, to assess the distribution of Nluc mRNA transcripts within the epithelial tissues of the lungs. The confocal images clearly showed the presence of Nluc transcripts throughout the lungs that were exposed to LNP/Nluc aerosol (Fig. 5B). In contrast, no signal was detected in the lungs exposed to PBS aerosol, validating the specificity of the assay (Fig. 5C). Moreover, the delivered Nluc mRNA transcripts were predominantly localized in the club cells, suggesting that the inhaled mRNA was effectively deposited in the lung epithelial tissues (Fig. 5C). These findings strongly support that our MAP produced aerosols fine enough to reach the lung epithelial cells of mice through spontaneous respiration, validating its efficacy in pulmonary mRNA delivery. To evaluate the effectiveness of the delivered mRNA in cell transfection, we conducted bioluminescence imaging on the collected lungs. Similarly, we aerosolized LNP/Nluc into the chamber, delivering a dose of 1 mg Nluc mRNA. After 24 hours post-inhalation, the mouse lungs were imaged *ex vivo*. The bioluminescence imaging showed consistent luciferase expressions in the collected lungs among all the mice. This consistent expression in the lungs supports the robust performance of our MAP in delivering mRNA to the lungs and achieving successful protein expression in the respiratory tissues of mice (Supplementary Fig. 11). We conducted further investigations to assess whether mRNA transfection was dose-dependent. The results demonstrated that the Nluc expressions in the lungs were directly proportional to the dose of LNP/Nluc aerosolized (Fig. 5D, E). This indicates that increasing the dose of LNP/mRNA led to a corresponding increase in the protein expression in the lungs. In addition, the site of mRNA transfection through inhaled LNP/Nluc was exclusively confined to the lungs (Fig. 5D). The conventional LNP formulation containing DLin-MC3-DMA exhibits inherent tropism for transfection to the liver when administered systemically. However, our results corroborate that inhaled LNPs can deliver mRNA to the respiratory system, consistent with previous studies^4^. We also conducted luminescence measurements at both 24 hours and 48 hours after treatment. It was shown that luciferase expression was higher at 24 hours post-treatment compared to 48 hours post-treatment (Fig. 5F, G). This observation indicates the transient nature of mRNA transfection, with the expression of the delivered mRNA reaching its peak at an earlier timepoint before gradually declining over time. Subsequently, we examined the possibility of acute lung damage following LNP/Nluc inhalation through histopathology. The mice were exposed to LNP/Nluc aerosol containing 1 mg of mRNA (Fig 5H) or an equivalent volume of PBS (Fig. 5I). After 24 hours of treatment, we collected the lungs, prepared lung sections, and stained them with hematoxylin and eosin (H&E) to assess any potential histological changes or tissue damage. In the H&E-stained lung sections, the lungs of both groups appeared histologically normal. However, in both groups, we observed minimal mononuclear cell infiltrates around the terminal bronchioles, interstitium, or peribronchiolar areas (Supplementary Fig. 12). Furthermore, the lungs showed a slight increase in lymphocytes in the bronchus-associated lymphoid tissues, but there was no significant difference between the two groups, suggesting that these abnormalities were likely artifactual findings and not related to the treatment. These minor changes could be attributed to the excessive accumulation of liquid in the tissues, resulting in mild congestion in the lungs. Overall, these findings collectively reinforce the pulmonary efficacy of our MAP, offering a safe, effective, and straightforward means of delivering mRNA therapies for respiratory conditions while avoiding unwanted effects in other organs.

## DISCUSSION AND CONCLUSIONS

Inhalable gene therapy holds tremendous potential for diverse drug development pursuits, including inhalable vaccines and therapies for inherited lung disorders like CF^4, 5, 8^. Even with recent discovery of selective organ-targeting LNPs leading to effective mRNA transfection of the lungs^9^, inhalation continues to stand out as the optimal, least invasive, and most effective approach to specifically reach the lung bronchioles and parenchyma while avoiding unintended systemic transfections. Given the expanding application of mRNA in gene editing^24, 25^, the interest in achieving highly accurate delivery through inhalation has grown substantially, particularly for its potential to offer a lasting solution to inherited lung disorders. However, ensuring a sufficient therapeutic dose remains a significant hurdle^26^. One plausible explanation for the limited pulmonary delivery could be the susceptibility of nanoparticles to destabilization during nebulization. Researchers have actively sought a solution to this deadlock, working on developing more robust nanoparticles through meticulous formulation adjustment^4, 15^ and exploration of new biomaterials^12, 13^. While these efforts are promising, the resolution of this challenge might ultimately stem from the innovation of advanced medical devices capable of generating aerosols while preserving the integrity of nanoparticles.

In this study, we have developed a new microfluidic system designed for producing nanoparticle aerosols suitable for inhaled mRNA therapy. Our findings affirm the capability of this MAP to generate a uniform nanoparticle aerosol without causing deformation in nanostructure or loss of encapsulated mRNA. Unlike conventional nebulizers such as vibrating mesh nebulizers and jet nebulizers, our MAP employs individually addressable nozzles for precise droplet ejection, allowing the generation of an aerosol containing LNP/mRNA at a lower operating frequency. Through this low shear aerosolization approach, our MAP prevents nanoparticle disruption, lipid aggregation, and mRNA leakage that occur with other nebulizers. Namely, this MAP maintains the size distribution and mRNA encapsulation of LNPs even after the aerosolization process. This was validated by cryoTEM imaging, which revealed minimal impact on LNP morphology compared to the distortion and dissociation observed with the mesh nebulizer. This preservation of nanoparticle integrity is critical not only for effective mRNA delivery to cells but also to prevent unwanted side effects in the respiratory system^27^. Furthermore, the MAP offers precise control over droplet size and plume dimensions, ensuring accurate dosing. Importantly, our device exhibited more efficient mRNA delivery efficiency to cells compared to the conventional nebulizer. While the mesh nebulizer led to a drastic 100-fold reduction in mRNA delivery, the MAP consistently maintained intracellular mRNA delivery efficiency. These results emphasize how the intactness of nanoparticles directly influence mRNA delivery efficiency. Lastly, despite the LNPs’ inherent tropism to the liver, the LNP aerosols generated by the MAP successfully delivered mRNA to the lungs of mice through inhalation. It was achieved in a dose-dependent manner, exhibiting selective lung transfection without causing significant inflammations. These findings highlight the potential utility of the platform for administering pulmonary mRNA therapies.

Recent advancements in LNP chemistry have identified the important roles of PEG molarity in RNA delivery. The incorporation of PEG into LNPs primarily serves to maintain the nanoparticles’ stability during self-assembly^1, 3^. In addition, PEG molecules hinder interactions between LNPs and serum proteins, extending the nanoparticles’ circulation time. However, they also inhibit the formation of a biomolecular corona on the nanoparticle surface, which delays the LNPs’ endocytosis and subsequent mRNA delivery^28^. In the context of LNP nebulization, PEG molecules are considered to contribute to the recovery or stabilization of nebulized nanoparticles through steric effects. This intricate interplay of PEG in LNP chemistry adds complexity to the formulation design for inhalation. Even with meticulous optimization, the nebulization process compromises LNP integrity, thus obscuring the key traits necessary for effective lung transfection^11^. We posit that our MAP has the potential to eliminate uncertainties in formulation discovery by preventing LNP deformation during nebulization. This platform holds the promise of enabling optimized LNP formulations, developed for systemic administration, to also perform effectively in inhalation scenarios. Alternatively, formulation screening based on intratracheal instillation could offer higher accuracy in predicting efficacies when delivered via aerosols^25^. Of note, the preservation of mRNA cargo integrity throughout the aerosolization process could facilitate precise dosing and reduce the risk of immunogenicity associated with activating RNA sensors in the patients’ lungs^29–31^. Collectively, we anticipate that our MAP will play a pivotal role in expediting the development of inhalable nanoparticles. Further, harnessing this platform has the potential to mark a significant leap in overcoming the challenges associated with translation of pulmonary gene therapy.

## MATERIALS AND METHODS

### Materials

Firefly luciferase (*Fluc*) mRNA was purchased from Trilink Biotechnologies. Cholesterol was obtained from Sigma-Aldrich (MO, USA), and DSPE-PEG_2K_ and DMG-PEG_2K_ were purchased from NOF America. DLin-MC3-DMA was purchased from Biofine International Inc. (BC, Canada). 1,2-distearoyl-sn-glycero-3-phosphocholine (DSPC), 1-palmitoyl-2-oleoyl-sn-glycero-3-phosphocholine (POPC), 1,2-dioleoyl-sn-glycero-3-phosphoethanolamine-N(lissamine rhodamine B sulfonyl) (Rhod-PE) and 1,2-dioleoyl-sn-glycero-3-phosphoethanolamine-N-(7-nitro-2-1,3-benzoxadiazol-4-yl) (NBD-PE) were obtained from Avanti Polar Lipids, Inc. (AL, USA).

### Characterization of droplets generated by a microfluidic aerosolization platform (MAP)

Liposomes were prepared as a substitute to LNPs in the characterization of droplet formation by a microfluidic aerosolization platform. Briefly, the lipid phase consisted of the following components: DSPC, cholesterol, and DSPE-PEG_2K_, in a 52:45:3 molar ratio, and was diluted to 25 mM in EtOH/DMF mixture (97.5:2.5). Sterile PBS and lipid solutions were heated at 65°C and mixed at a 2:1 volumetric ratio using a NanoAssemblr Benchtop system (Precision Nanosystems, BC, Canada), followed by overnight dialysis against sterile PBS at 4°C in 10 kDa Slide-a-Lyzer G2 cassettes (Thermo Fisher, MA, USA). The batch size of liposomes was 1.2 L in total. Hydrodynamic size and polydispersity of the liposomes were determined with DLS in a Zetasizer Nano ZSP device (Malvern Panalytical, UK).

Droplet formation and ejection was evaluated using the drop watcher system (JetXpert Dropwatcher, ImageXpert, NH). Droplets that were dispensed by the microfluidic head were examined, and the behavior of a single drop was captured using the drop watcher camera. The system was calibrated to a ratio of 1 pixel to 0.001034 mm using internal software and a calibration target provided by the manufacturer. A known-width slit was imaged at a determined magnification and working distance. It allows for the determination of droplet sizes and the estimation of droplet volumes using the software provided by the manufacturer.

Droplet size was measured using Spraytec^®^ droplet size measurement system (Malvern Panalytical, UK). The system measures size distribution of sprayed droplets via their laser diffractions. This requires the angular intensity of light scattered from a spray to be measured as it passes through a laser beam. The recorded scattering pattern is analyzed and plotted using the manufacturer’s software.

The thermal simulation was conducted using a proprietary FMS ejector modeling code (Funai Lexington, KY) with the following settings: bubble detachment time of 1236 ns, voltage of 11V, chip temperature of 45°C, pre-pulse duration of 200 ns, pulse delay of 800 ns, and main heat-pulse duration of 600 ns. A proprietary aqueous dye formulation with well-defined thermophysical properties was used to characterize the fluid dynamics of the MAP. Data were plotted using Origin 2022 (ver. 9.9).

### *In vitro* transcription of mRNA

*In vitro* transcription of mRNA was performed as described^32^. Briefly, a linearized plasmid containing nanoluciferase (*Nluc*) under a T7 promoter was used as a template for *in vitro* transcription. *Nluc* mRNA was synthesized using the HiScribe T7 high yield RNA synthesis kit (New England Biolabs Inc., MA, USA) with CleanCap Reagent AG (TriLink Biotechnologies) according to the manufacturer’s instructions. Synthesized mRNA was purified using the Monarch RNA cleanup kit (New England Biolabs) and stored at −80°C. Concentration of *Nluc* mRNA was measured using a multimode microplate reader (Tecan Trading AG, Switzerland). To visualize the mRNA and assess for degradation, agarose gel electrophoresis was performed. 1 µg of IVT mRNA or RiboRuler high range RNA ladder (Thermo Fisher, MA, USA) was denatured and loaded on 1.5% agarose-formaldehyde gel prestained with GelRed (Biotium, CA, USA). The gel was run at 85 V for 2 h, followed by UV visualization.

### LNP Formulation

LNPs were prepared using microfluidic mixing as previously described^11^. Lipid phase consisted of the following components: DLin-MC3-DMA, cholesterol, DMG-PEG_2K_, and DSPC in a 50:38.5:1.5:10 molar ratio and were diluted to 5.5 mM in 100% ethanol. mRNA was diluted in sterile 50 mM citrate buffer. The mRNA and lipid solutions were mixed at a 3:1 volumetric ratio using a NanoAssemblr Benchtop system (Precision Nanosystems, BC, Canada), followed by overnight dialysis against sterile PBS at 4°C in 10 kDa Slide-a-Lyzer G2 cassettes (Thermo Fisher, MA, USA).

### LNP Characterization

Hydrodynamic size and polydispersity of the LNPs were determined with DLS in a Zetasizer Nano ZSP device (Malvern Panalytical, UK). Encapsulation of mRNA was quantitatively determined by a modified protocol using Quant-iT RiboGreen RNA assay kit (Thermo Fisher, MA, USA) and a multimode microplate reader. Encapsulation of mRNA was measured qualitatively before and after aerosolization using agarose gel migration.

### LNP Aerosolization

To aerosolize LNPs through ultrasonication, LNPs were added to a vibrating mesh nebulizer (Aerogen, Ireland) dropwise as previously described^4^. For microfluidics-based aerosolization with the microfluidic device, LNPs were added to a single channel of the previously described microfluidic cartridge, connected to the MAP (FMS, Lexington, KY). Droplet ejection was controlled using FMS-provided software with the following settings: frequency of 1.2 kHz, target temperature of 25°C, voltage of 11V, preheat pulse conditioning of 100 ns, preheat pulse duration of 200 ns, main-heat pulse conditioning of 800 ns, and main-heat pulse duration of 600 ns.

### FRET-based lipid membrane fusion assay

Lipid mixing and membrane fusion induced by shear stress during aerosolization were assessed using a FRET assay^22, 23^. Briefly, DOPE-conjugated FRET probes, NBD-PE and Rho-PE, were incorporated into FRET liposomes. This formulation causes a reduction in NBD fluorescence due to FRET to rhodamine. When lipid fusion occurred, an increase in NBD signal would be detected owing to the greater distance between the two probes. To prepare FRET liposomes, DOPC:NBD-PE:Rho-PE mixture was combined in chloroform at a molar ratio of 99:0.5:0.5 in a round-bottom flask. The chloroform was eliminated through constant airflow and flask rotation, resulting in a uniform lipid firm on the flask bottom. Vacuum was applied for 2 h to ensure complete chloroform evaporation. After drying, the lipid film was sonicated for 20 min, hydrated with HEPES-buffered saline, and extruded using a mini extruder (Avanti Polar Lipids). The liposome solution was passed through a 100 nm membrane 11 times. Subsequently, FRET liposomes were mixed with LNP/mRNA at a 1:1 (v/v) ratio and exposed to different conditions: no treatment, 2% Triton X-100, 20-min bath sonication, vibrating mesh nebulizer, or MAP. The resulting samples were added to a black 96-well plate (80 µl per well) for fluorescence measurement. Fluorescence measurement (*F*) was conducted on a multimode microplate reader at Ex/Em = 465/535 nm. The results from no treatment and Triton X-100 treatment were set as negative (*F*_*min*_) and positive control (*F*_*max*_), respectively. Fusion% was calculated as (*F* - *F*_*min*_)/(*F*_*max*_ - *F*_*min*_) x 100.

### Cryogenic Transmission Electron Microscopy (CryoTEM)

Following plasma cleaning of grids, 2 µl of sample was deposited onto the grid in the FEI Vitrobot chamber at 100% relative humidity and allowed to rest for 10 seconds. The grid was then blotted for 1 second with filter paper and subsequently plunged into liquid ethane cooled by liquid nitrogen. The frozen grids were inspected for visible defects and then intact grids were assembled into cassettes. CryoTEM acquisition was performed with a Glacios cryoelectron microscope equipped with a Gatan K3 camera at 200 kV.

### Cell Culture

HeLa cells were gifted from Prof. Robert Langer at Massachusetts Institute of Technology. A549 cells were kindly provided by Prof. Adam Alani at Oregon State University. 16HBE14o-cells and CFBE41o- cells were from Prof. Kelvin MacDonald at OHSU. HEK293T/17 were purchased from American Type Culture Collection (ATCC). HeLa and HEK293T/17 cells were cultured in DMEM supplemented with 10% heat inactivated FBS, 1% penicillin/ streptomycin, and 10mM HEPES buffer. 16HBE14o- cells and CFBE41o- cells were maintained in MEM/EBSS supplemented with 10% heat inactivated FBS, 1% penicillin/ streptomycin/ glutamine, 1% sodium pyruvate, and 10mM HEPES buffer. A549 cells were cultured with RPMI-1640 supplemented with 10% heat-inactivated FBS, 1% penicillin/ streptomycin, and 10mM HEPES buffer.

### Characterization of *in vitro* mRNA delivery

Cells were seeded on a white 96 well plate at 4,000 cells/ well, on a clear 12 well plate at 50,000 cells per well, or on an 8 well chamber microslide (Ibidi) at 40,000 cells per chamber, followed by overnight incubation for attachment. Aerosolization of mRNA was conducted with one of two methods: a vibrating mesh nebulizer (Aerogen, Ireland) and a MAP (Funai Lexington, KY). Cells in particular plates were treated with corresponding doses of mRNA. For cells in the 96 well plate, nebulized particles were collected from output of nebulizing unit and added to seeded cells. For cells in the 12 well plate, nebulizing unit was held and dispensed mRNA encapsulating LNPs directly over cells, followed by a 24 h incubation. Cell viability and *in vitro* luciferase expression was evaluated via ONE-Glo + Tox luciferase reporter and cell viability assay kit (Promega) and a multimode microplate reader.

### Animals

All animals studied were conducted at Oregon Health and Sciences University and approved by the Institutional Animal Care and Use Committee (IACUC, IP0001707).

### *In vivo* LNP/mRNA delivery through microfluidic aerosolization

For characterizing mRNA delivery to mice through aerosolized LNPs, a whole chamber aerosolization system was chosen as described^12^. Briefly, all mice of group were placed in a 3 L chamber at once. LNPs encapsulating mRNA were prepared at 0.25 mg/mL, diluted in PBS. LNPs were loaded into a microfluidic cartridge single channel at a volume of 200 µl at a time. LNPs were aerosolized into the chamber at a rate of 25 µl per 2 minutes until the total dose of mRNA 1 mg total delivered. All mice in group were unrestrained without sedation. After 24 h post-treatment, the animals were sacrificed, and the lungs were collected for further characterization. To detect Nluc expression in the lungs, the collected lungs were gently and briefly washed in sterile PBS, incubated in 200 µl of 40-fold dilution of Nano-Glo substrate (Promega) in PBS per lung at room temperature for 5 minutes. *Ex vivo* bioluminescent imaging was then conducted using the IVIS Lumina XRMS (PerkinElmer). Collected lungs were then homogenized by adding the lungs to vials, adding 150 µl PBS to each lung and completely homogenizing with a hand-held homogenizer over ice until a slurry is observed. Samples moved to microcentrifuge tubes were centrifuged at 17,000 xg for 30 minutes at 4°C. On a white 96 well plate, 30 µl of the supernatant and 60 µl of 80- fold dilution of Nano-Glo substrate in PBS were incubated at room temperature for 5 minutes. Luminescence was read with a multimode microplate reader. Luminescence was normalized with total protein concentration, which was measured with a BCA protein assay kit (Thermo Fisher).

### Histopathology

BALB/c mice were exposed to LNP/mRNA aerosols generated from the microfluidic device until the total dose of mRNA 1 mg total delivered or sterile PBS until the equal volume delivered. After 24 h, mice were humanely euthanized, and their lungs were perfused with sterile PBS through the right ventricle. A 20G catheter was inserted into the trachea for lung inflation using a 10% neutral buffered formalin solution under a pressure of 25 cm from the surgical plane. The inflated lungs were carefully removed, placed in tissue embedding cassettes, and immersed in 70% ethanol for the purpose of dehydration. Following this, the tissues were embedded in paraffin, sectioned, placed onto slides, stained using H&E, and coverslipped, enabling histopathological assessment (IDEXX BioAnalytics, MO, USA). Whole-slide imaging of the specimens was conducted using a slide scanner (Leica Biosystems), and the images were analyzed using Aperio ImageScope v12.4.6.5003 (Leica Biosystems) and Fiji.

### RNAscope *in situ* hybridization

Delivered Nluc mRNA and endogenous scgb1a1 mRNA transcripts that were present in formalin-fixed paraffin-embedded (FFPE) lung tissue sections were visualized using RNAscope Multiplex Fluorescent Reagent Kit v2 (ACD) according to the manufacturer’s protocol. Nluc probe (Cat. No. 885981), and Scgb1a1 probes (Cat. No. 420351-C3) were prepared, and tyramide signal amplification (TSA)-based Opal fluorophores, Opal 570 (Akoya Biosciences, #FP1488001KT, 1:800 dilution) and Opal 690 (#FP1487001KT, 1:1,500 dilution), were used to visualize Nluc and Scgb1a1 transcripts. Confocal images were obtained with a ZEISS LSM 880 (Carl Zeiss AG).

## AUTHOR INFORMATION

G.S. conceived the idea. J.K., A.J., and G.S. designed experiments and analyzed the data. J.K., A.J., E.B., and N.M. conducted characterization of aerosolized nanoparticles. J.K., A.J., and E.B. designed and performed mouse inhalation experiments. Y.E. performed and interpreted cryoTEM imaging. M.M. and B.J. fabricated a microfluidic platform and characterized fluid properties. The manuscript was written by J.K., A.J., and G.S. with contributions of all authors. All authors have given approval to the final version of the manuscript.

## ACKNOWLEDGMENT

We would like to thank Dr. Manish Giri for coordination in experiments and assistance in manuscript review and feedback. This project was supported through funding from the National Heart Lung and Blood Institute (NHLBI) 1R01HL146736-01 (G.S) and an SRA from Rare Air Health, Inc. (G.S).

