## Supplemental File for "Microfluidic platform enables shear-less aerosolization of lipid nanoparticles for messenger RNA inhalation"

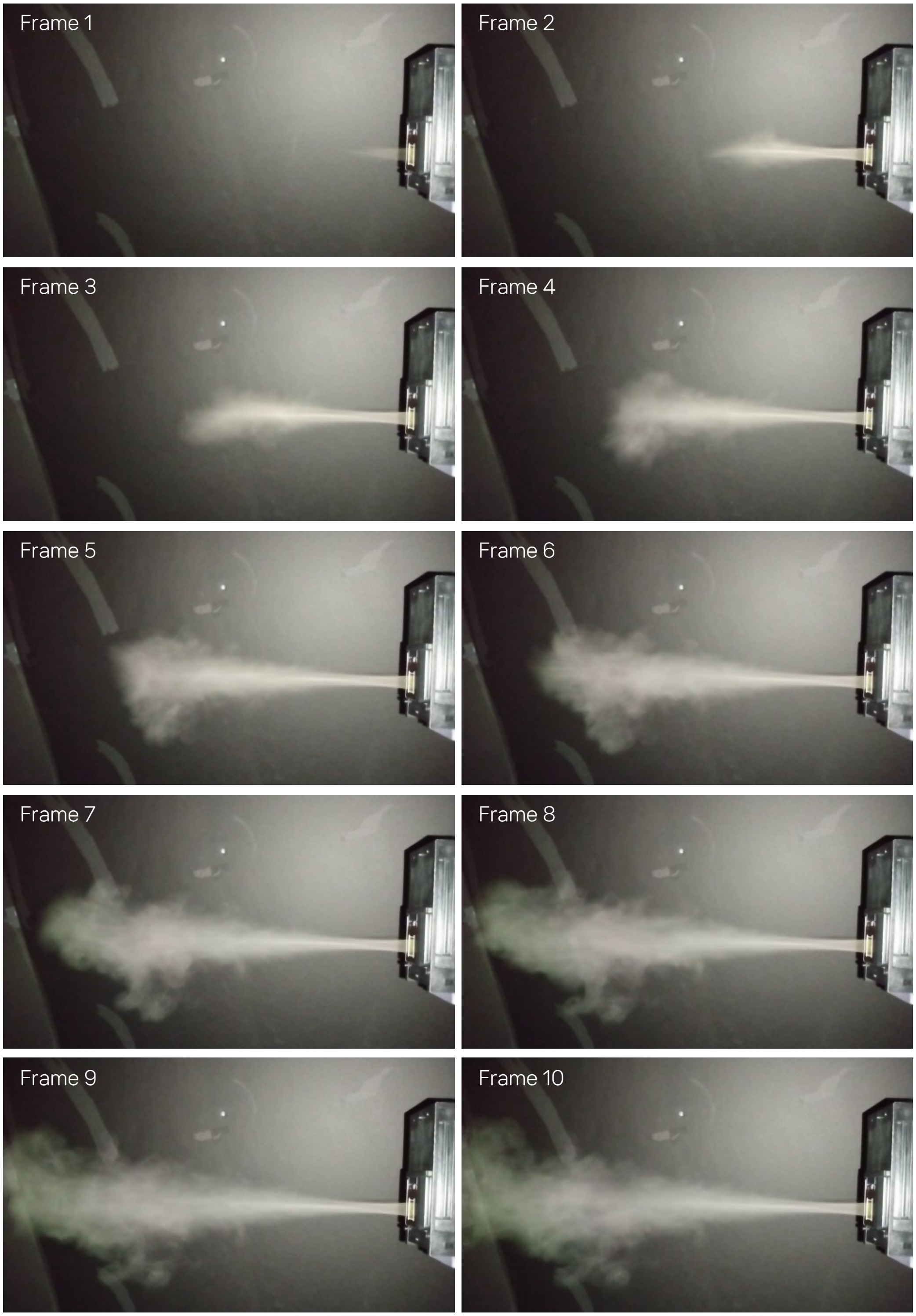


**Supplementary Fig. 1. An image sequence of aerosol generation from the microfluidic device.**

**Cumulant Results**

| | |
|---|---|
| **Z-Avg (nm):** | 81.03 |
| **Pd Index:** | 0.084 |
| **Pd (nm):** | 23.5 |
| **%Pd:** | 29.0 |
| **Derived kcps:** | 2616.9 |

**Distribution Results**

| | Size (d.nm): | % Int | σ | %Pd |
|---|---|---|---|---|
| **Peak 1:** | 89.17 | 100.0 | 28.57 | 32.0 |
| **Peak 2:** | 0.000 | 0.0 | 0.000 | 0 |
| **Peak 3:** | 0.000 | 0.0 | 0.000 | 0 |

**Undersize Results**

| Di (%) | Size (d.nm): |
|---|---|
| 50 | 84.7 |
| 90 | 132 |
| 95 | 146 |

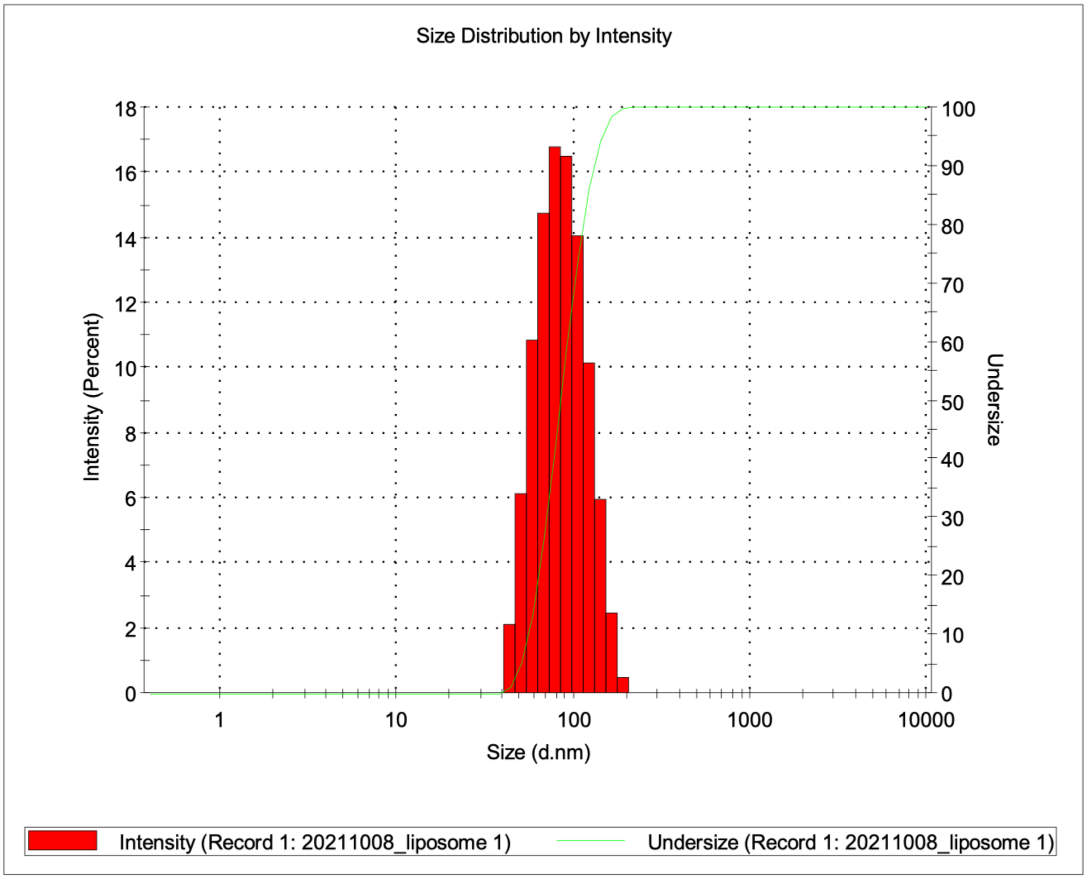


**Supplementary Fig. 2. A representative size distribution of liposomes.**

(A)

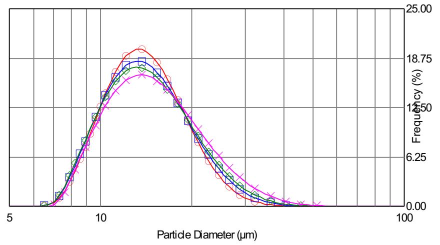


| | File | Dx(10) | Dx(50) | Dx(90) | Lot |
|---|---|---|---|---|---|
| [N] | 20 Full | 9.65 | 13.80 | 20.76 | ... |
| [N] | 30 Full | 9.49 | 13.88 | 21.75 | ... |
| [N] | 40 Full | 9.45 | 14.00 | 22.52 | ... |
| [N] | 60 Full | 9.68 | 14.58 | 24.36 | ... |

[V]=Volume [N]=Number

(B)

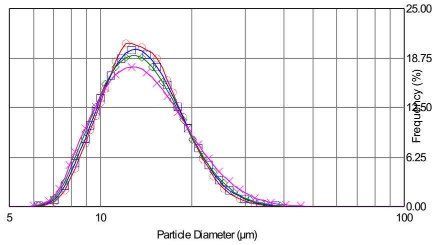


| | File | Dx(10) | Dx(50) | Dx(90) | Lot |
|---|---|---|---|---|---|
| [N] | 20 Spatial | 9.54 | 13.50 | 19.88 | ... |
| [N] | 30 Spatial | 9.39 | 13.46 | 20.23 | ... |
| [N] | 40 Spatial | 9.24 | 13.35 | 20.48 | ... |
| [N] | 60 Spatial | 9.09 | 13.44 | 21.60 | ... |

[V]=Volume [N]=Number

(C)

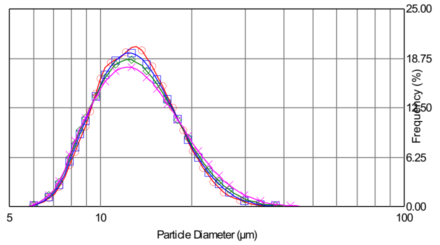


| | File | Dx(10) | Dx(50) | Dx(90) | Lot |
|---|---|---|---|---|---|
| [N] | 20 Temporal | 9.05 | 12.89 | 19.19 | ... |
| [N] | 30 Temporal | 8.94 | 12.87 | 19.59 | ... |
| [N] | 40 Temporal | 8.82 | 12.87 | 20.04 | ... |
| [N] | 60 Temporal | 8.79 | 12.99 | 20.91 | ... |

[V]=Volume [N]=Number

**Supplementary Fig. 3. Measuring the size distribution of aerosol droplets at various distances from the device.** Laser scattering was measured at various distances from the nozzle plates: 20 mm (red), 30 mm (blue), 40 mm (green), and 60 mm (magenta). Aerosolization was conducted from (A) all nozzles at 15 kHz (full), (B) half of the total nozzles at 15 kHz (spatial), or (C) all nozzles at 7.5 kHz (temporal).

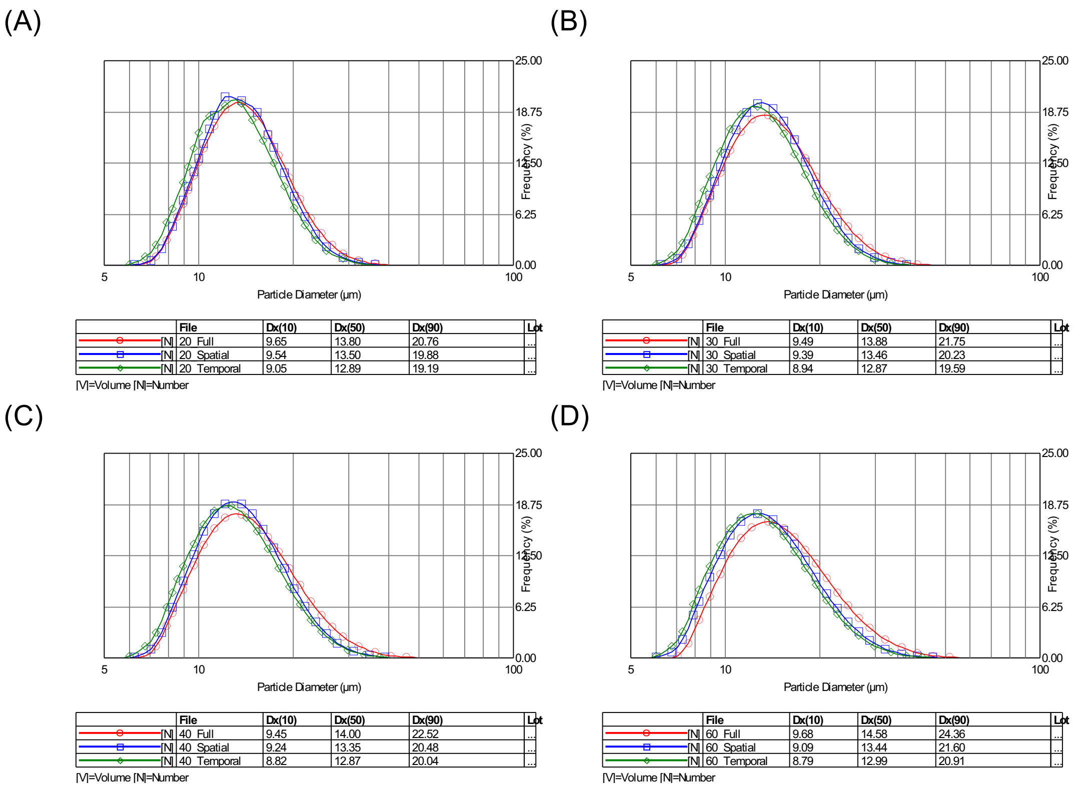


**Supplementary Fig. 4. Measuring the size distribution of aerosol droplets by varying aerosolization frequencies or the number of nozzles.** Aerosolization was conducted from all available nozzles at 15 kHz (red; full), half of the total nozzles at 15 kHz (blue; spatial), or all nozzles at 7.5 kHz (green; temporal). Laser scattering was measured at various distances from the nozzle plates: (A) 20 mm, (B) 30 mm, (C) 40 mm, and (D) 60 mm.

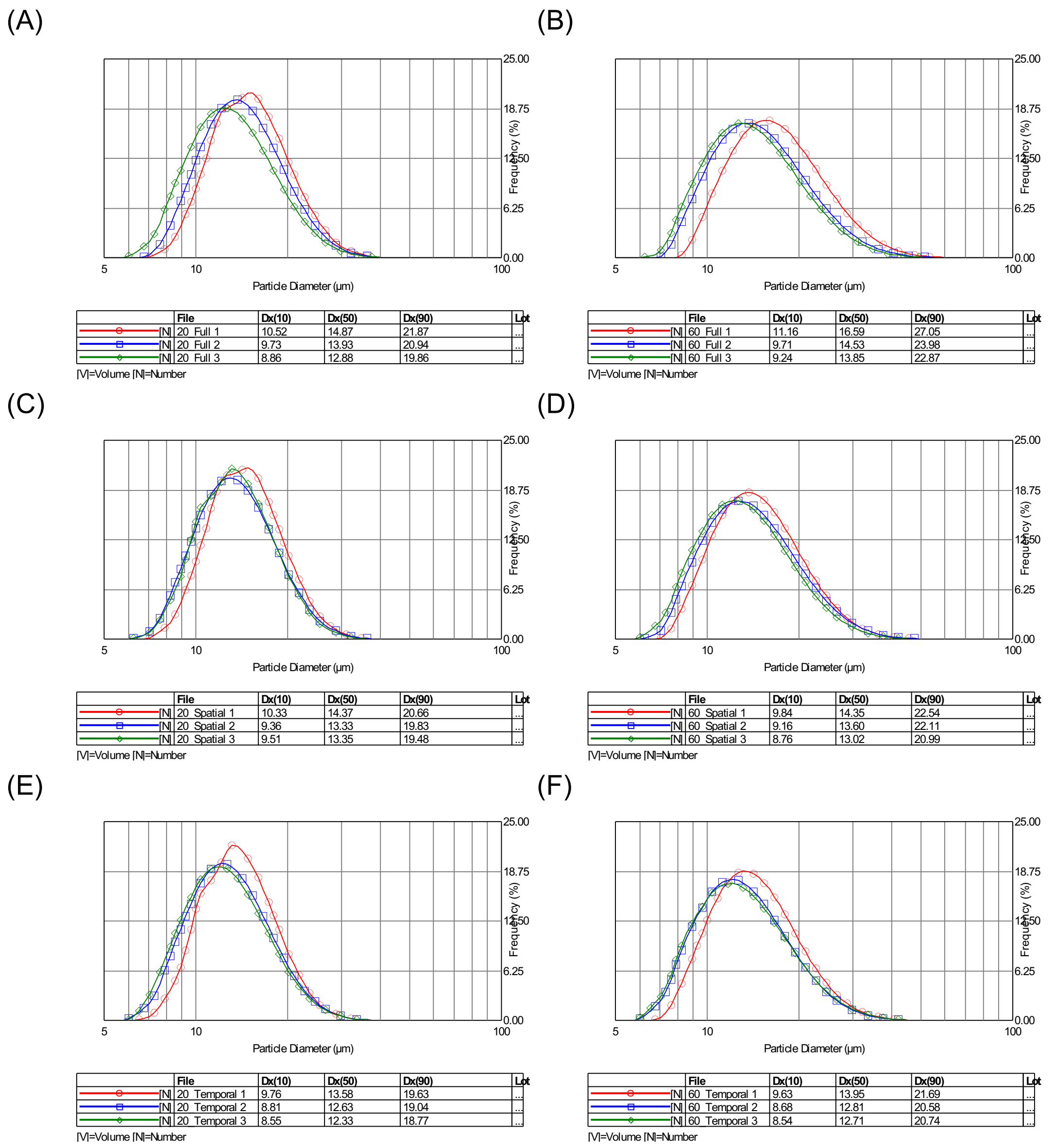


**Supplementary Fig. 5. Measuring the size distribution of aerosol droplets at various elapsed times of aerosolization.** Laser scattering was measured at (A, C, E) 20 mm or (B, D, F) 60 mm from the nozzle plates. Measurements were recorded in the beginning (red), middle (blue), or end (green) of the dispense. Aerosolization was conducted from (A, B) all available nozzles at 15 kHz (full), (C, D) half of the total nozzles at 15 kHz (spatial), or (E, F) all nozzles at 7.5 kHz (temporal).

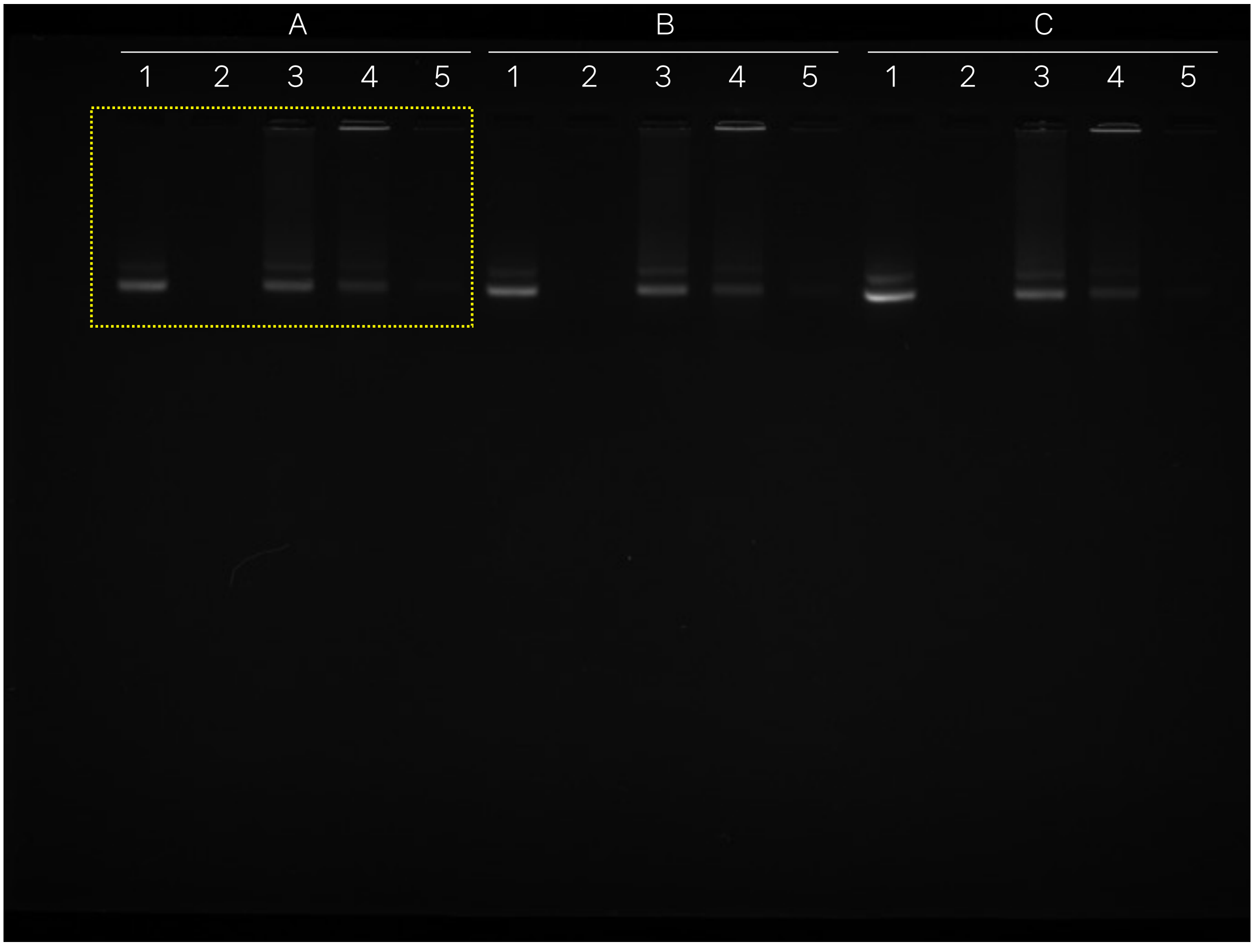


| # | Samples |
|---|---|
| 1 | mRNA only |
| 2 | LNP before treatment |
| 3 | LNP + Triton X-100 |
| 4 | LNP after vibrating mesh |
| 5 | LNP after microfluidic device |

**Supplementary Fig. 6. Agarose gel electrophoresis analysis of LNP encapsulation of mRNA after mesh nebulizer or MAP.** (A-C) Replicate samples: 1) mRNA only, 2) LNP before treatment, 3) LNP treated with Triton X-100, 4) LNP after nebulization with a vibrating mesh, and 5) LNP after microfluidic device. Yellow dotted line indicates gel image used in Fig. 3G.

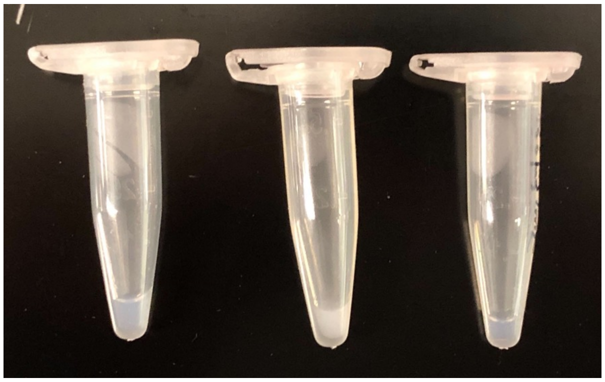


1. LNP Solution
2. Vibrating Mesh
3. Microfluidic

**Supplementary Fig. 7. Visual change in opacity of LNP solution after aerosolization.** LNP solutions before (1) and after vibrating mesh (2) or microfluidic aerosolization (3) were imaged to show changes in opacity in the solutions.

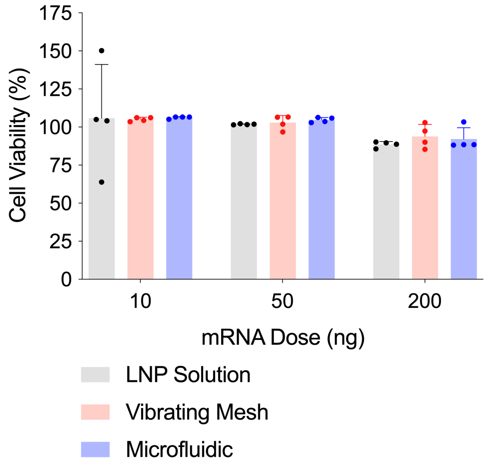


**Supplementary Fig. 8. Cell viability in response to LNP transfection with aerosolized LNPs.** Cell viability of 293T/17 cells treated with LNP/Fluc solution (grey), LNP/Fluc aerosolized by a vibrating mesh nebulizer (red) or the microfluidic platform (blue) at various mRNA doses. (n=4).

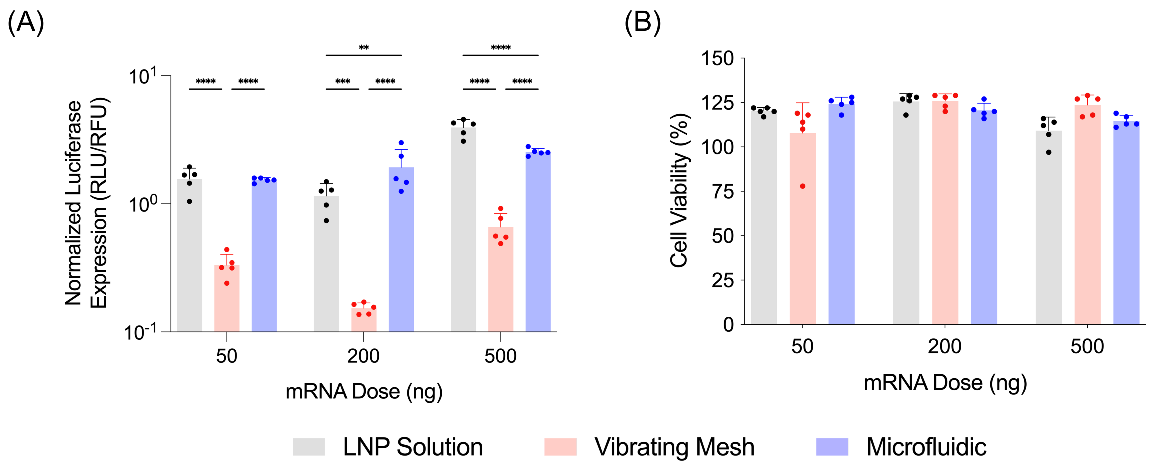


**Supplementary Fig. 9. Aerosolized LNP delivery to relevant lung cell line. (**A) Normalized luciferase expression and (B) cell viability of human bronchial epithelium (16HBE14o-) cells treated with LNP/Fluc solution (grey), LNP/Fluc aerosolized by a vibrating mesh nebulizer (red) or the microfluidic platform (blue) at various mRNA doses. (n=5). *$p<0.05$; ***$p<0.001$; ****$p<0.0001$.

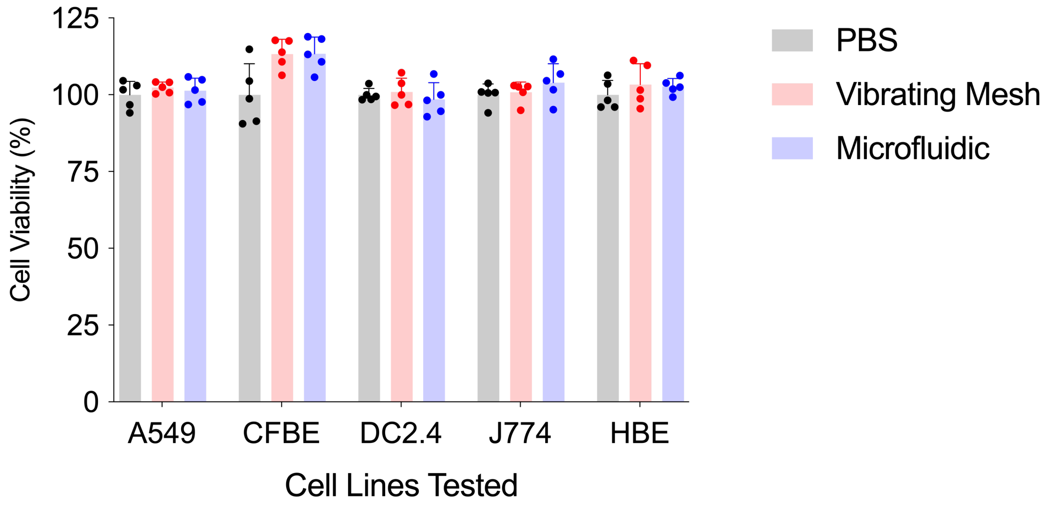


**Supplementary Fig. 10. Cell viability of various cell lines after treatment with aerosolized LNPs.** Cell viability of various cell lines treated with PBS (grey), LNP/Fluc aerosolized by a vibrating mesh nebulizer (red) or the microfluidic platform (blue) at a dose of 50 ng FLuc mRNA per well (n=5). CFBE: CFBE41o- human bronchial epithelial cells, J774: mouse macrophage cells, and HBE: human bronchial epithelium 16HBE14o- cells.

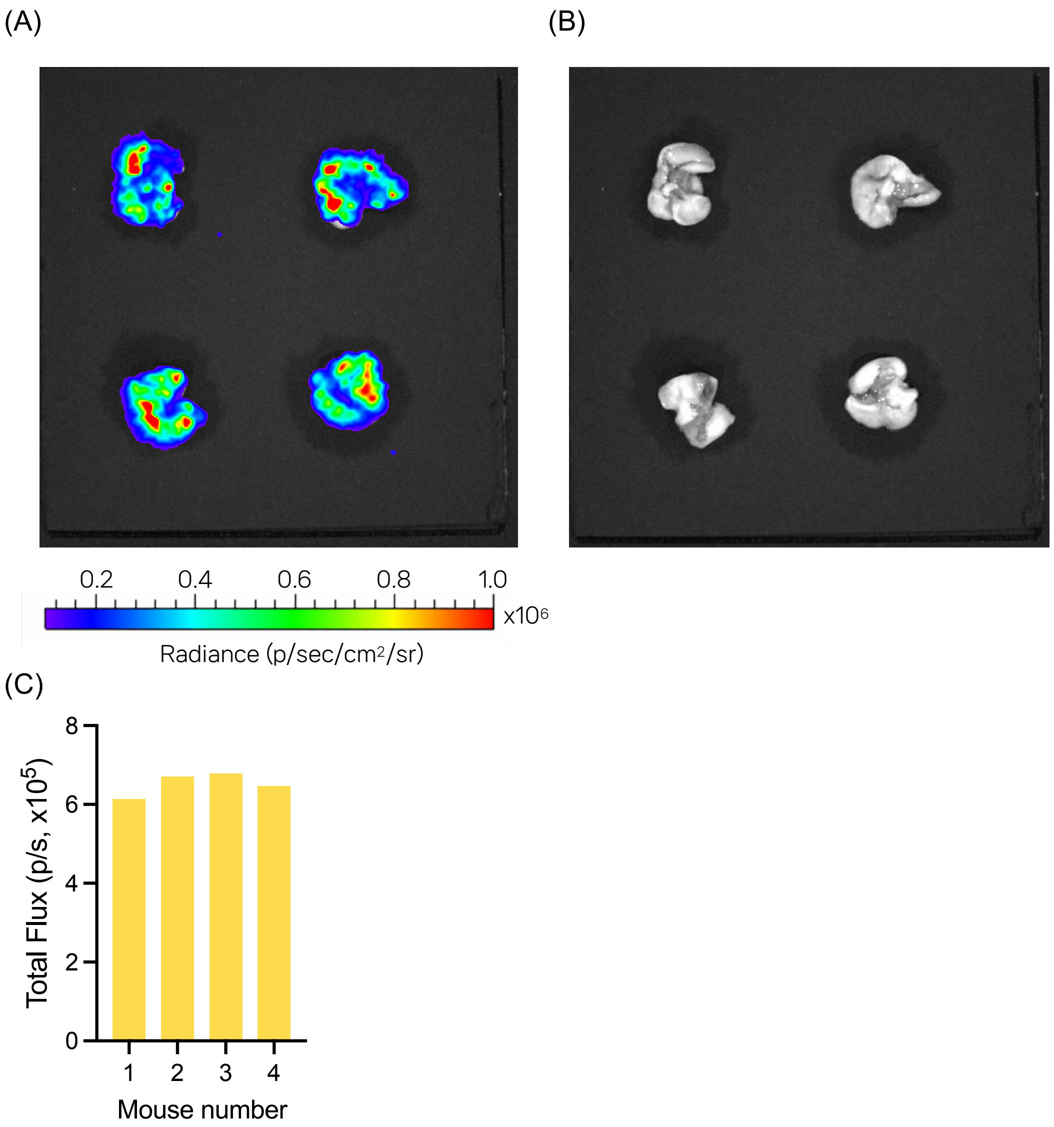


**Supplementary Fig. 11. Lung delivery of aerosolized LNP/Nluc by spontaneous inhalation.** (A-C) Microfluidic device delivery consistency across all 4 mice in a whole-body rodent inhalation system. (A) Bioluminescent signals and (B) a photograph of the collected lungs. (C) Quantified luminescent signals in the captured images.

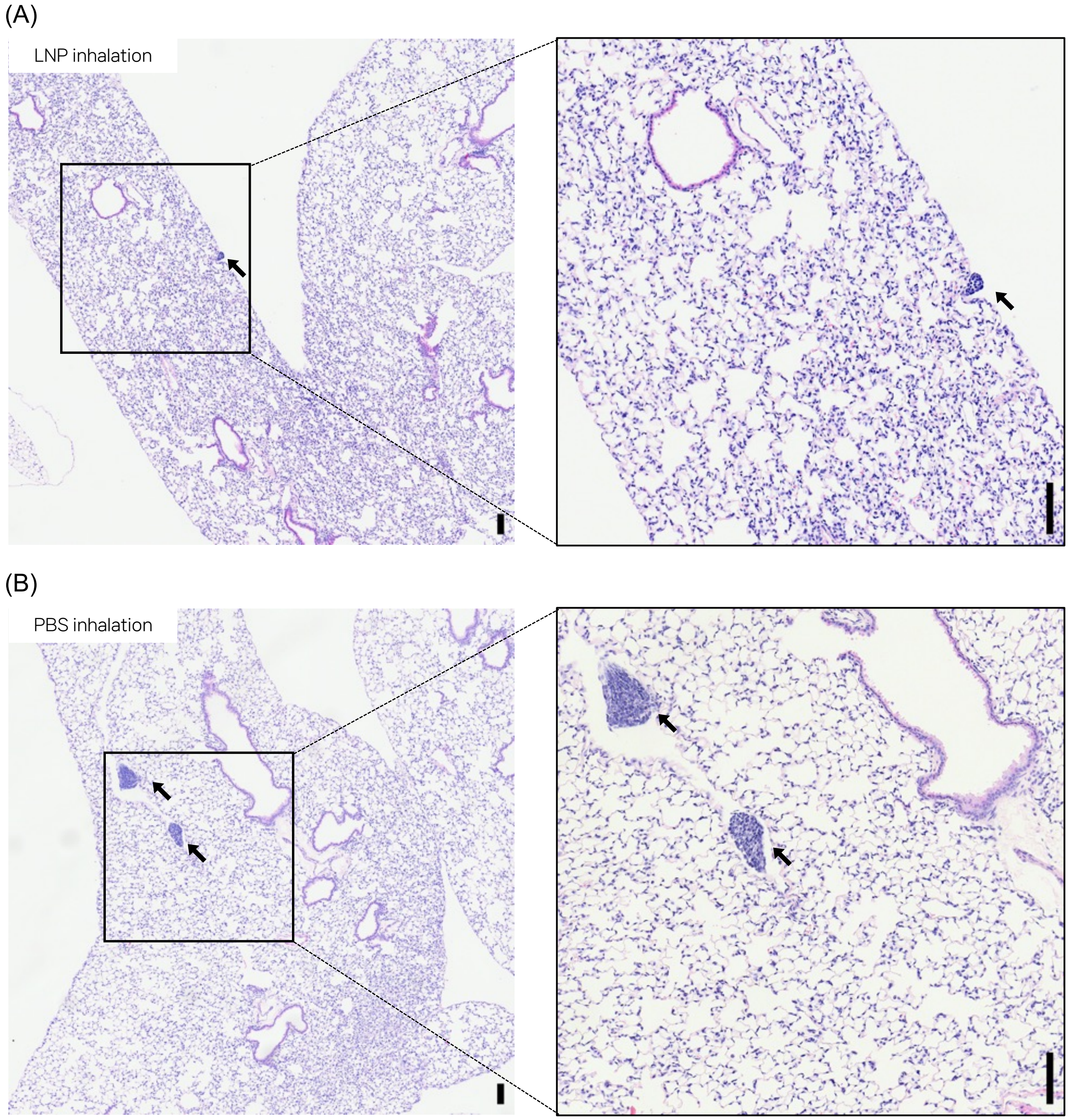


**Supplementary Fig. 12. Histopathological analysis of mouse lungs collected 24 h after (A) inhalation of LNP/Nluc when 1 mg of mRNA was aerosolized, or (I) inhalation of an equal volume of sterile PBS using the microfluidic platform.** Insets indicate the area of interest. Arrows indicate minimal increases of lymphocytes in the bronchus-associated lymphoid tissue (BALT). Scale bars indicate 100 $\mu$m.
